# Stress alters the activity of Leydig cells dependently on the diurnal time

**DOI:** 10.1101/2021.02.18.431858

**Authors:** Marija Lj Medar, Silvana A Andric, Tatjana S Kostic

**Author notes:** **Correspondence:** Tatjana S Kostic, Dositeja Obradovica Sq 2, Novi Sad, 21000, Serbia.

## Abstract

**Background:** The increasing amount of data points to the circadian timing system as an essential part of processes regulating androgen homeostasis. However, the relationship between stress response, timekeeping-, and steroidogenesis-related systems is unexplored.

**Objective:** The purpose of the study was to analyze the stress-response of the testosterone-producing Leydig cells depending on the stressful event’s time.

**Materials and methods:** The study was designed to follow the effects of 3-hour immobilization (IMO) applied at different periods during the day. The IMO performed once (1xIMO) or repeated in 10 consecutive days (10xIMO). Principal-component-analysis (PCA) followed the expression study of the clock and steroidogenic-related genes in Leydig cells.

**Results:** Both types of IMO in all investigated periods increased corticosterone and decreased testosterone blood level. Transcriptional analysis revealed different sensitivity to IMO events depending on the circadian time. The majority of steroidogenesis-related genes (*Lhcgr, Cyp11a1, Cyp17a1, Hsd3b1/2*) were down-regulated in the inactive but unchanged or even up-regulated in the active phase of the day. Both types of IMO potentiated the expression of clock elements *Bmal1*/BMAL1, *Per1/*PER1 regardless of the day’s stage and reduced *Reverba* in the inactive phase. The PCA confirmed a major shift, for both IMO-types, in the transcription of steroidogenesis and clock genes across the inactive/active phase. Further, the diurnal pattern of the glucocorticoid receptor (*Nr3c1*/GR) expression in Leydig cells was increased in the inactive phase due to 10xIMO. The observed time-dependent IMO-response of the Leydig cells correlated with different GR engagements.

**Discussion:** Stress- and the circadian-system coordinatively shape Leydig cell’s physiology, assuming diverse GR engagement as a possible factor in mediating the diurnal effect of stress.

**Conclusion:** The Leydig cell’s stress-response depends on the time of the stressful situation, emphasizing the importance of circadian activity in supporting androgen homeostasis and male fertility.

## Introduction

The increase in evidence points to the circadian timing system as an essential part of processes regulating reproduction. The circadian timing system provides an organism with the sense of time of day to ensure that physiological and behavioral events coincide to give offspring ^1^. Temporal coordination of different organs is mediated by the actions of the master clock in hypothalamic suprachiasmatic nuclei, together with hormonal and nervous systems employing multi-level signaling through negative and positive feedback loops. The content of hormonal signals is overwhelmingly conveyed in a rhythmic secretory pattern. In cells, including cells of the reproductive tract, the timing system consists of clock genes coding transcription factors that operate through positive (BMAL1, CLOCK, NPAS2, RORA,B) and negative (PER1-3, CRY1,2, REV-ERA,B, DEC) elements organized in transcription-translation regulatory feedback loops ^2,3^. The rhythm in clock gene transcription is decoded into physiological signals through cell-specific transcriptional activity via circadian promoter elements such as E-boxes, D-boxes, and ROREs ^4^.

In males, testosterone production is a rhythmic process predominantly regulated by the rhythm of pulsatile LH secretion and clock elements ^5,6^. Those cells rhythmically express clock genes (*Bmal1, Per1-3, Cry1,2, Reverba,b, Dec, Rora,b*) as well as major steroidogenic genes (*Star, Cyp11a1*, and *Cyp17a1*) followed by an oscillatory pattern of blood testosterone ^7,8^. In humans and rats, testosterone reaches a peak at the beginning of the active phase ^7–9^. Testosterone is a crucial hormone regulating male reproduction, including the development and maintenance of the male phenotype; consequently, potential changes in circadian rhythmicity of testosterone production/secretion could impact reproductive function.

It is known that stress disturbs male reproduction by decreasing blood testosterone ^10–12^ acting on all three levels of the hypothalamic-pituitary-testicular axis ^13^. The effect is achieved through the neural and hormonal system, possibly associated with central and peripheral clocks ^14^.

In addition, some earlier studies suggest stress-response dependence on time of stress event ^15,16^, pointing to the relationship between the circadian rhythm of life functions and the stress-response. However, molecular events and signaling pathways responsible for such differences are not described.

Keeping in mind the importance of sustainable circadian rhythm on homeostasis, including maintaining reproductive ability, this study aims to explore the relationship between stress response and circadian timekeeping in testosterone-producing Leydig cells. Precisely, this study estimated the stress-response of the Leydig cells dependently on the time of the stressful event.

## Materials and methods

### Ethical Approval

All the experiments were carried out in the Laboratory for Chronobiology and Aging, the Laboratory for Reproductive Endocrinology and Signaling, DBE, Faculty of Sciences at the University of Novi Sad. All experiments were approved by the Ethical Committee on Animal Care and Use of the University of Novi Sad (statement no. 01-201/3), operating under the rules of the National Council for Animal Welfare and following statements of National Law for Animal Welfare (copyright March 2009). All experiments were performed and conducted by following the European Convention for the Protection of Vertebrate Animals used for Experimental and other Scientific Purposes’ (Council of Europe No 123, Strasbourg 1985) and NIH Guide for the Care and Use of Laboratory Animals (NIH Publications, No. 80 23, revised, 7th ed., 1996). All experiments were in adherence to the ARRIVE guidelines.

### Animals & stress model

Adult (age, 3 mo; body wt, 250–270 g) male *Wistar* rats, bred and raised in the Animal Facility of the Department of Biology and Ecology (Faculty of Sciences, University of Novi Sad), were used for the experiments. The rats were raised in controlled environmental conditions (22 ± 2°C; 14h light:10h dark; lights on at 6:00 AM) with food and water ad libitum. The chosen experimental model of stress was psychophysical stress induced by 3h immobilization (IMO). IMO was performed as described previously ^10,17^. Briefly, rats were bound in the supine position for 3h by fixing their limbs to the wooden board with thread. The head motion was not limited. At the end of experiments, rats were quickly decapitated without anesthesia. All activities during the dark phase were performed under the red light.

The two types of experiments were performed depending on number od IMO session: (1) to analyze circadian sensitivity on acute IMO (1xIMO), rats were subject to IMO in different periods during 24h (ZT0-3, ZT8-11, ZT14-17, and ZT20-23; ZT0 is a time when the light turned on); (2) to examine the effect of repeated IMO, rats were immobilized each day at the same time (ZT0-3, ZT8-11, and ZT20-23) for 10 consecutive days (10xIMO). Corresponding groups of freely moving rats served as controls in both IMO experiments.

For both tipes of IMO experiments, rats for control and experimental groups were randomly divided into 4 or 3 time points consisted each from 5-6 rats per time points. No masking was used during group allocation, data collection or data analysis. Previous results from our group obtained on the rat model showed that the size of the group of 5-6 rats was sufficient to calculate hormone parameters of the circadian rhythm ^7,8^. The sample size was checked also by Power Analysis using G Power software (http://core.ecu.edu/psyc/wuenschk/Power.htm) according to previous results. Experiments were repeated two times.

### Preparation of purified Leydig cells

The testes were isolated, decapsulated, and the main blood vessel was removed as it was previously described (Kostic et al., 2010; Baburski et al., 2016, 2015). Decapsulated testes were placed in plastic tubes (2 testes per tube) containing collagenase solution (0.25 mg collagenase/mL-1.5 %-BSA-20 mM HEPES-M199, Sigma, St Louise, Missouri) and incubated in a shacking-water bath (15 min/34 °C/120 cycles/min). The reaction was stopped by adding a sufficient amount of cold M199 medium, and seminiferous tubules were removed by filtration. The interstitial cells’ suspension was centrifuged (160xg/5 min), and the cell pellet was resuspended in DMEM-F12 medium (Sigma, St Louise, Missouri) 8 mL/rat. The resuspended cells (35-40×10^6^) were moved on the Percoll gradients (Sigma, St Louise, Missouri) and centrifuged 1100xg/28 min. After centrifugation, Leydig cells were harvested, washed with a sufficient amount of the M199-0.1% BSA, centrifuged (200xg/5 min), and resuspended in DMEM-F12 medium (5 mL/animal). In line with the procedure and according to the HSD3B staining ^20^ presences of Leydig cells in the culture was greater than 90%; according to the Trypan blue exclusion test ^7^, viability of the cells was greater than 90%. The Leydig cell suspension was centrifuged, and the pellet was stored at -80 °C for RNA and protein analysis.

### RNA isolation and cDNA synthesis

Total RNA from purified Leydig cells was isolated using the using Trizol (TermoFisher Scientific, Waltham, USA) or Rneasy kit reagent following the protocol recommended by the manufacturer (QIAGEN, Valencia, California). The amount of total RNA’s and purity were assesed by spectrophotometric measurement (NanoDrop, Shimadzu). After DNase I treatment, cDNA was synthesized using the Applied Biosystems kit (Termo Fisher Scientific, Massachusetts, USA) according to the manufacturer’s instructions. Analyses of appropriate and relevant positive and negative controls were performed for both DNase treatment and cDNA synthesis.

### Real-time PCR and relative quantification

Relative gene expression was done by RQ-PCR using SYBR Green-based chemistry (Applied Biosystems, Foster City, California) in the presence of a 5 μl aliquot of the reverse transcription reaction product (25 ng RNA calculated on the starting RNA) and specific primers. The primer sequences used for RQ-PCR analysis and GenBank accession codes for full gene sequences (www.ncbi.nlm.nih.gov/sites/entrez) are provided in Suppl. Table 1. *Gapdh* was also measured in the same samples and used to correct variations in RNA content. The relative quantification of each gene was performed in duplicate.

### Protein extraction and western blot analysis

Platted cells were lysed by buffer (20 mM HEPES, 10 mM EDTA, 2.5 mM MgCl_2_, 1 mM DTT, 40 mM β-glicerophosphate, 1% NP-40, 2 µM leupeptin, 1 µM aprotinin, 0.5 mM AEBSF, phosphatase inhibitor cocktail (PhosSTOP, Roche, Basel, Switzerland) and the lysates were uniformed by Bradford method. Samples were mixed with SDS protein gel loading solution (Quality Biological, Gaithersburg, USA), boiled for 5 min, and separated by one-dimensional SDS-PAGE electrophoresis (Bio-rad, Berkeley, USA).

Transfer of the proteins to the PVDF membrane was performed by electroblotting overnight at 4°C/40A, and the efficacy of transfer was checked by staining and distaining the gels. The membranes were blocked by 3% BSA-1xTBS for 2h/room temperature, incubation with primary antibody was done overnight at 4 °C, and 0.1% Tween-1xTBS was used for washing membranes incubation with secondary antibody was done for 1h/room temperature ^6^. Immunoreactive bands were detected using luminol reaction and MyECL imager (www.Fischer.sci)/films and densitometric measurements were performed by Image J program version 1.32 (https://rsbweb.nich.gov//ij/download.html). Anti-BMAL1 purchased from Abcam (Cambridge, UK, #ab49421, dilution 1:800); anti-PER1 purchased from Abcam (Cambridge, UK, #ab3443, dilution 1:500); anti-GR purchased from Thermo Fisher Scientific (Waltham, Massachusetts, USA, #MA1-510, dilution 1:1000); anti-ACT purchased from Santa Cruz Biotechnology (Heidelberg, Germany, #sc-8432, dilution 1:800). All antibodies used in this study are validated and previously published by our group ^6,7,21^.

### Hormone and glucose measurements

Androgen level was measured by RIA and it was referred to testosterone + dihydro-testosterone (T + DHT) because the anti-testosterone serum No. 250 showed 100% cross-reactivity with DHT ^18,19^. All samples were measured in duplicate in one assay (sensitivity: 6 pg per tube; intraassay coefficient of variation 5–8%). For serum corticosterone levels ^22^, all samples were measured in duplicate, in one assay by the corticosterone EIA Kit (Caymanchem, Michigan, USA) with 30 pg/ml as the lowest standard significantly different from blank. The level of glucose in serum was estimated by using BAYER Contour plus Glucose Monitoring System (Bayer, Basel, Switzerland; measuring range 0.6-33.3 mmol/L). Technically, all serum samples obtained individually, were measured in duplicate.

### Statistical and rhythm analysis

The results represent group means ± SEM values of two in vivo experiments. Rhythm parameters (p, MESOR, amplitude, and acrophase) were obtained by cosinor.lm() function from the “cosinor” package ^23^ in the R environment and by using online Cosinor.Online software fitted to 24 h period (http://www.circadian.org/softwar.html) ^24^. Statistical significance (p < 0.05) between the treated group and control group within the same time point and between the groups including all time points (p < 0.05) were analyzed using Mann–Whitney test and two-way ANOVA, respectively.

Principal component analysis (PCA) was done with dudi.PCA() function implemented in “ade4” package ^25^, on scaled and centered data matrix, within the R environment. We decided to retain the first two PC based on eigen values and cumulative variation. In support of such a decision, we performed Horn’s parallel analysis for a PCA with the “paran” package, to adjust for finite sample bias in retaining components ^26^. Biplots visualization were done with “factoextra” package ^27^. The effects of 1xIMO and 10xIMO on gene expression were presented by using The Clustered Image Map (CIM) analysis (https://discover.nci.nih.gov/cimminer/oneMatrix.do).

## Results

### Time-dependent effect of IMO on hormone levels

To examine whether the circadian time of stress event can provoke different responses, rats were immobilized for 3h, in different periods during 24h, once (1xIMO) or repeatedly for 10 days (10xIMO), and responses were analyzed immediately after the IMO session.

As expected ^21^, corticosterone, glucose, and testosterone show diurnal pattern in the serum of control rats (Fig. 1A&B&C; the rhythm parameters in Suppl. Table 2). The peak of these oscillations occurs around ZT11 for all three. 1xIMO effectively increased corticosterone (Fig. 1A) and glucose (Fig. 1B) levels in all investigated ZT, although glucose levels were increased in the beginning of inactive and at the end of active phase (ZT3 and ZT23). On the contrary, 1xIMO reduced testosterone levels in serum in all investigated ZTs (Fig. 1C).

**Figure 1.**
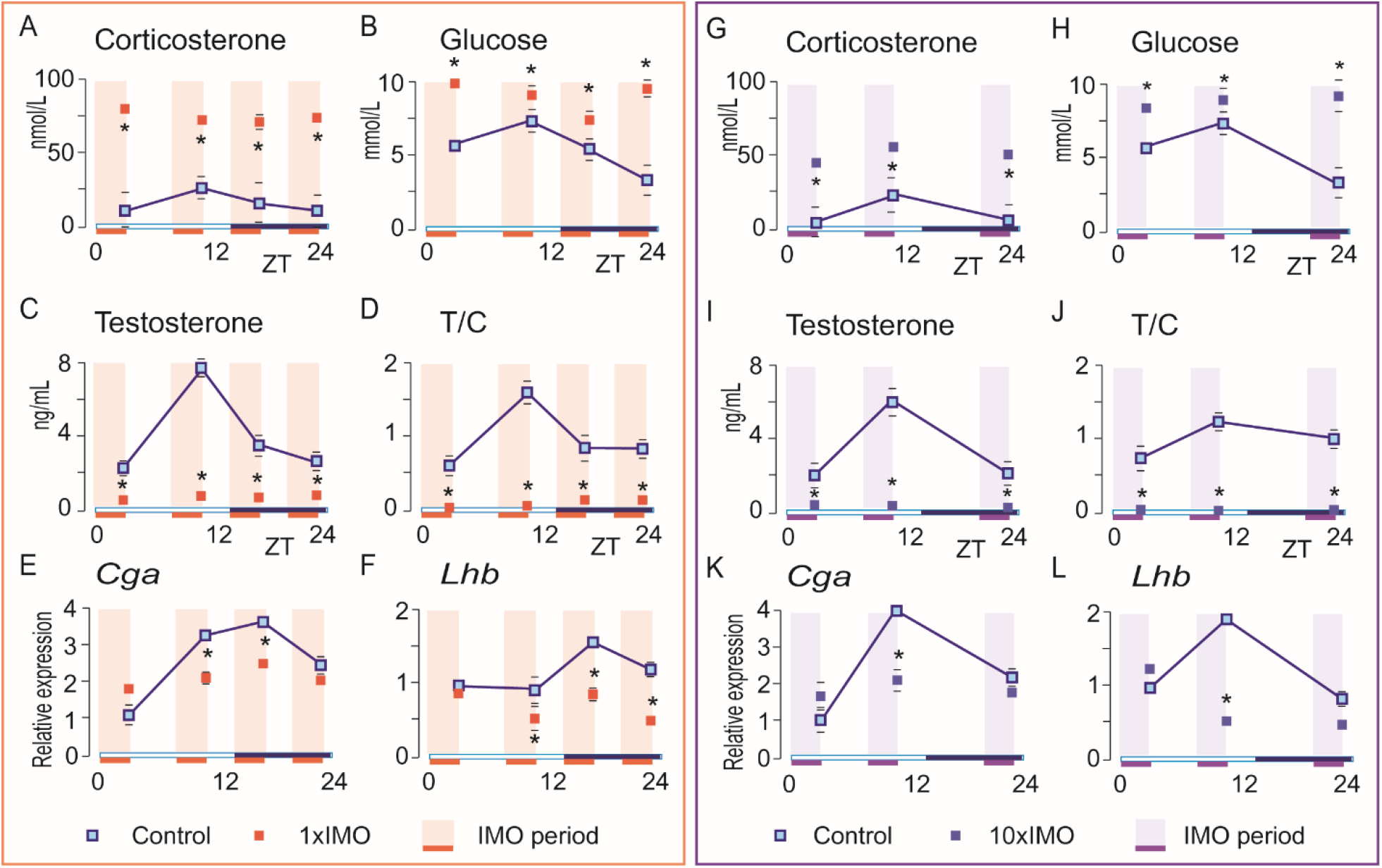
Time-dependent effect of IMO on hormone levels and pituitaries gene levels. Wistar rats were subject to 1xIMO for three hours in four-time points (ZT0-3, ZT8-11, ZT14-17, ZT20-24) or to 10xIMO in three-time points (ZT0-3, ZT8-11, ZT20-24) during the 24h. Serum was collected for determination of corticosterone, glucose, and testosterone levels as well as testosterone/corticosterone (T/C) ratio (1xIMO: A-D; 10xIMO: G-J). Pituitaries were collected, RNA isolated for RT followed by RQ-PCR analysis of genes encoding the LH subunits (1xIMO: E, F; 10xIMO: K, L). Data points are group mean ± SEM values, (n = 10-12 for A, B, C, G, H, I, J, and n=5-6 for E, F, K, L). For possible rhythm prediction please see Suppl. Table 2. ^*^Statistical significance between control and IMO groups for the same time point (p < 0.05).

Analysis of the relationship between blood testosterone and corticosterone (T/C) could point to the ability of the Leydig cells to produce testosterone in stress conditions. In that respect, the relative T/C was calculated on ZT3 in control group (ZT3=1). Obtained results depicts circadian fluctuations in blood testosterone and corticosterone; the peak of fluctuations observed in the control group was at ZT11 (T/C ZT11=1.6). Nevertheless, 1xIMO reduced T/C approximately 38-fold, depending on time of IMO event. The lowest T/C value was observed in ZT3 (T/CZT3 = 0.02), while the highest was observed in ZT11 (T/CZT11 = 0.03) (Fig. 1D).

The repetition of stress events over ten days always at the same time, 10xIMO, increased blood corticosterone (Fig. 1G), and glucose (Fig. 1H). However, increments of corticosterone were one-third lower than after 1xIMO at all tested times (Fig. 1G). In addition, the repeated IMO profoundly reduced testosterone (Fig.1I), which resulted in a substantially decreased T/C ratio (Fig. 1J). Described patterns were observed in all examined ZTs.

Since LH is the principal regulator of Leydig cell endocrine function, the transcriptional pattern of the genes encoding pituitary LH subunits (*Cga, Lhb*) were studied. Both genes showed a diurnal transcriptional model in control conditions (Fig.1 E, F; please see rhythm parameters in Suppl.Table2). 1xIMO significantly decreased *Cga* at ZT11 and ZT17 (Fig. 1E) and *Lhb* at ZT11, ZT17, and ZT23 (Fig. 1F). The 10xIMO turn-down transcription of *Cga* (Fig. 1K) and *Lhb* (Fig. 1L) at ZT11.

### The time-dependent effects of IMO on the steroidogenesis-related genes expression in Leydig cells

The endocrine function of the Leydig cell entirely depends on the expression pattern of the steroidogenic elements. Therefore, to detect a possible different time-dependent stress effect on the gene expression essential for the Leydig cells’ endocrine function, we perform transcriptional analysis in Leydig cells originating from control and IMO rats in various periods during 24h.

The *Lhcgr*, encoding LH receptor, did not show a diurnal pattern of transcription, (Fig. 2A). 1xIMO significantly decreased the *Lhcgr* transcription in the light phase, while the dark phase was without effect (Fig. 2 A).

**Figure 2.**
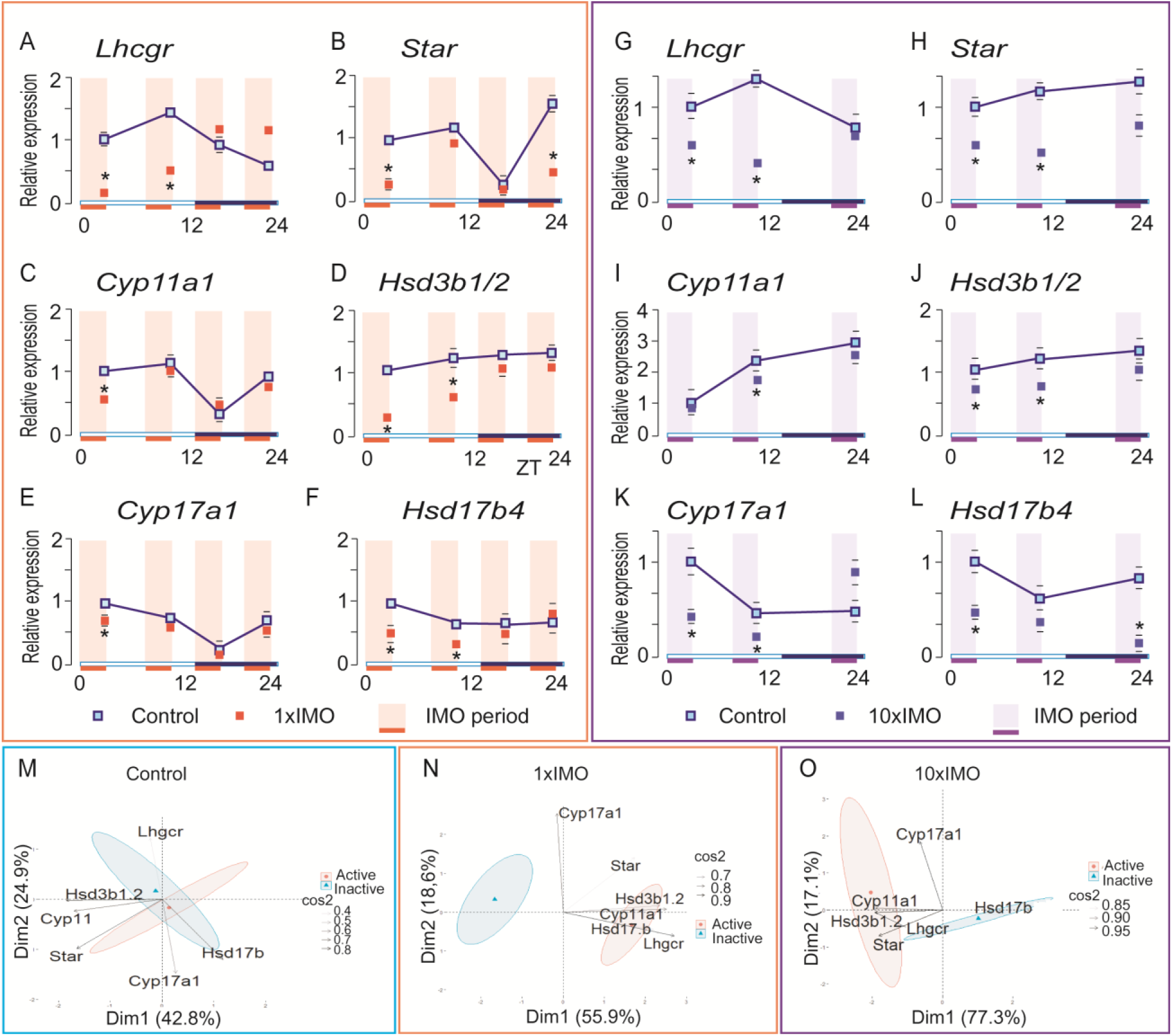
The time-dependent effects of IMO on the expression of the steroidogenesis-related genes in Leydig cells. *Wistar* rats were subject to 1xIMO for three hours in four time points (ZT0-3, ZT8-11, ZT14-17, ZT20-24) or to 10xIMO in three time points (ZT0-3, ZT8-11, ZT20-24) during the 24h. Leydig cells were purified, RNA extracted and used for RT followed by RQ-PCR analysis of steroidogenesis-related genes expression (1xIMO: A-F; 10xIMO: G-L). Data points are group mean ± SEM values, (n = 5-6). For possible rhythm prediction please see Suppl. Table 2. ^*^Statistical significance between control and IMO groups for the same time point (p < 0.05). PCA of steroidogenic-related genes depending on active/inactive phase in control (M), 1xIMO (N) and 10xIMO (O) conditions; Dim1 and Dim2 represents the first two PC and % of retained variation. cos2 estimates the qualitative representation of variables. In M, N and O datasets were prepared as a value that deviates from control at a given time (Supplemental table 3, 4 and 5).

Key steroidogenic genes, *Star, Cyp11a1, Cyp17a4*, exhibit a diurnal transcriptional pattern which reach a maximum approximately in the middle of the light phase (Fig. 2B, C, E, Suppl.Table2), a few hours earlier than the peak of blood testosterone. 1xIMO changed the expression pattern of the *Star, Cyp11a1*, and *Cyp17a4*, underlying a pronounced reduction in the inactive (light) phase (Fig. 2B, C, E). Although, results did not support diurnal pattern in the transcription of *Hsd3b1/2* and *Hsd17b4*, 1xIMO significantly reduced transcription of both genes during the inactive (light) phase (Fig. 2D, F).

The time-dependent effects of stress response were also seen in 10xIMO conditions (Fig. 2 G-L). The *Lhcgr* (Fig. 2G), *Star* (Fig. 2H), *Cyp11a1* (Fig. 2I), *Hsd3b1/2* (Fig. 2J), and *Cyp17a4* (Fig. 2K) were down-regulated in inactive without significant effect in active period. In addition, the *Hsd17b4* (Fig. 2L) was down-regulated by 10xIMO in all three examined ZTs.

Thus, the results showed different expression of steroidogenic genes over 24 h, including different IMO effects on their expression: in control samples, inactive and active phases tend to separate across the first two PC for 67.7% of the total dataset of steroidogenesis-related genes variability (Fig. 2M; variable loadings are shown in Suppl. Table 3); in 1xIMO and 10xIMO PC accounts for 74.5% (Fig. 2N, variable loadings are shown in Suppl. Table 4) and 94.4% (Fig. 2O, variable loadings are shown in Suppl. Table 5) data variability, respectively. In 1xIMO conditions, inactive and active phases are strongly driven by distinct expression profiles of *Lhgcr, Hsd3b1/2, Cyp11a1*, and *Hsd17b4* (Fig. 2N). Similarly, in 10xIMO conditions the transcription of *Lhgcr, Hsd3b1/2, Cyp11a1* and *Star* were less changed in the active than in the inactive phase (Fig. 2O). Such gene expression profiles point to the robustness of steroidogenesis-related genes (light and dark phase, respectively) under different IMO treatments, which may pinpoint them as a landmark for stress conditions.

### The time-dependent effects of IMO on the expression of the clock genes in Leydig cells

The effect of IMO on the peripheral clock in Leydig cells was estimated by transcription analysis of canonical clock genes (Fig. 3; Suppl.Table2).

**Figure 3.**
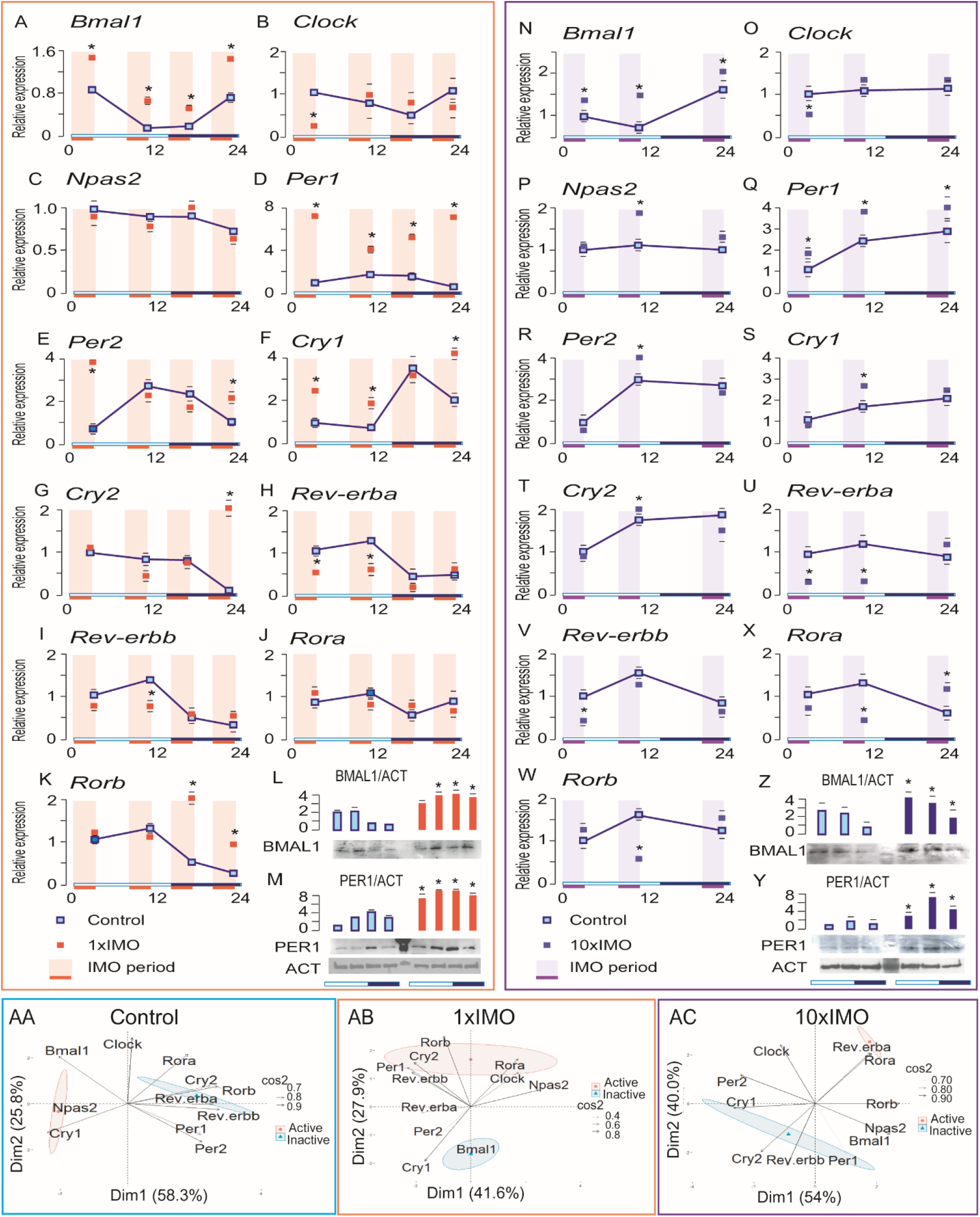
The time-dependent effects of IMO on the expression of the clock-related genes in Leydig cells. Wistar rats were subject to 1xIMO for three hours in four-time points (ZT0-3, ZT8-11, ZT14-17, ZT20-24) or to 10xIMO in three-time points (ZT0-3, ZT8-11, ZT20-24) during the 24h. Leydig cells were purified, RNA extracted, and used for RT followed by RQ-PCR analysis of steroidogenesis-related genes expression (1xIMO: A-K; 10xIMO: N-W). Data points are group mean ± SEM values, (n = 5-6). For possible rhythm prediction please see Suppl. Table 2. The protein level of BMAL1 (1xIMO: L; 10xIMO: Z) and PER1 (1xIMO: M; 10xIMO: Y) were determined by western blot. Representative blots are shown. Bars above blots represent mean ± SEM values from pooled data using scanning densitometry values normalized on ACT. ^*^Statistical significance between Control and IMO group at the same time point (p < 0.05). Expression pattern of clock genes depending on active/inactive phase in Leydig cells from control (AA), 1xIMO (AB) and 10xIMO (AC) rats analized by PCA; Dim1 and Dim2 represents the first two PC and % of retained variation. cos2 estimates the qualitative representation of variables. In AA, AB and AC datasets were prepared as a value that deviates from control at a given time (Supplemental table 6,7 and 8).

The majority of examined genes (*Bmal1, Per1/2, Cry1/2, Rev-erba/b, Rob*) had a diurnal expression in rat Leydig cells (Fig. 3; Suppl.Table2). On the contrary, *Clock, Npas*, and *Rora* transcripts did not follow any significant pattern (Fig. 3 B, C, J; Supplement Table 2). *Bmal1* transcript was the biggest at the beginning (Fig. 3A) while *Per1/2* (Fig. 3D, E), *Cry2* (Fig. 3G), *Rev-erba/b* (Fig. 3H, I), *and Rorb* (Fig. 3K) at the end of the light phase, followed by *Cry1* (Fig. 3F) rise in the dark phase. Generally, 1xIMO changed transcription of clock genes in Leydig cells: *Bmal1* increased in all IMO-times (Fig. 3A), and *Clock* (Fig. 3B) decreased immediately after morning IMO (ZT3) while its counterpart *Npas2* was unaffected by IMO situations (Fig. 3C). However, clock negative regulators in the circadian clock’s primary loop were enhanced: *Per1* after all IMO sessions regardless of the time of occurrence (Fig. 3D), and *Per2* at ZT3 and ZT23, *Cry1* at ZT3, ZT11 and ZT23, *and Cry2* at ZT23 (Fig. 3E, F, G). Additionally, IMO affected the transcription of genes in the secondary loop by decreased expression of *Rev-erba/b* during the inactive phase (Fig. 3H, I) and *Rorb* at ZT17 and ZT23 (Fig. 3K). Changed protein levels of BMAL1 and PER1 due to IMO were confirmed by western blot analysis. As expected, BMAL1 and PER1 showed circadian profiles in the control samples with an increased level of BMAL1 in the light in relation to the dark phase and vice versa situation for its negative regulator PER1 (Fig. 3L, M). IMO increased BMAL1 (Fig. 3L) and PER1 in all examined time points (Fig. 3M).

10xIMO altered clock genes expression in a similar way as 1xIMO (Fig. 3N-W): the *Bma1* increased and (Fig. 3N) *Per1* increased in all tested IMO-periods (Fig. 3Q) while *Reverba* decreased by IMO-sessions in light phase (Fig. 3U). 10xIMO efficiently increased BMAL1 (Fig. 3Z) and PER1 (Fig. 3Y) after all analyzed time-points.

Likewise previous, a PCA was run to convince into IMO effects on diurnal fluctuations of clock genes expression in Leydig cells. As expected, the diurnal phase transition is characterized by a distinct expression pattern of clock genes, *Rev-erbb/a, Rora/b, Cry2* in the inactive (light) phase, and *Cry1* during the active (dark) phase (Fig. 3AA, variable loadings are shown in Suppl. Table 6). The *Bmal1* and *Per1/2* had opposite directions (Fig. 3A, variable loadings are shown in Suppl. Table 6). Additionally, PCA confirmed that light and dark phases have different expression patterns under 1xIMO or 10xIMO conditions. In 1xIMO conditions, a pronounced cluster of genes in active (*Rorb, Cry2, Rev-erbb, Per1)*, opposes *Bmal1* in inactive phase and contributes to the separation between phases (Fig. 3AB, variable loadings are shown in Suppl. Table 7). Interestingly, *Clock, Npas2, Per2* but also *Rev-erba* are minorly contributing to the separation between phases (Fig. 3AB, variable loadings are shown in Suppl. Table 7). Additionally, 10xIMO treatment drives a robust difference in expression of *Rev-erba* and *Rora* during the active and *Rev-erbb* and *Cry2*, during inactive phase (Fig. 3AC, variable loadings are shown in Suppl. Table 8). Also, the light and dark phase shows the most distinctive change/shift across all analyzed data (94.0% variability), which implies that 10xIMO is strongly affecting the clock genes expression profiles (Fig. 3AC, variable loadings are shown in Suppl. Table 8).

### The time-dependent effects of IMO on the *Nr3c1*/GR expression in Leydig cells

Glucocorticoids are the primary hormonal mediators of stress with a large regulatory impact on Leydig cell’s steroidogenesis as well as survival ^10^. In that respect, the time-dependent *Nr3c1*/GR expression was monitored in Leydig cells from control and 1xIMO as well as 10xIMO rats. In Leydig cells from control rats, the diurnal expression pattern of *Nr3c1* and GR was observed (Fig. 4) with increased expression at the end of the light and the beginning of the dark phase. That 24h expression pattern of GR correlates with blood corticosterone fluctuation (R=0.885), allowing efficient regulation of target genes in Leydig cells. However, 1xIMO in the rest period reduced *Nr3c1* transcript (Fig. 4A) without significant effect on GR (Fig. 4B). Still, a decrease in *Nr3c1* was not detected in the case of 1xIMO in the active period. On the contrary, IMO repetition (10xIMO), in rest period, elevated *Nr3c1* (Fig. 4C) and GR (Fig. 4D) in Leydig cells. Again, the effect of stress was less pronounced in the active period.

**Figure 4.**
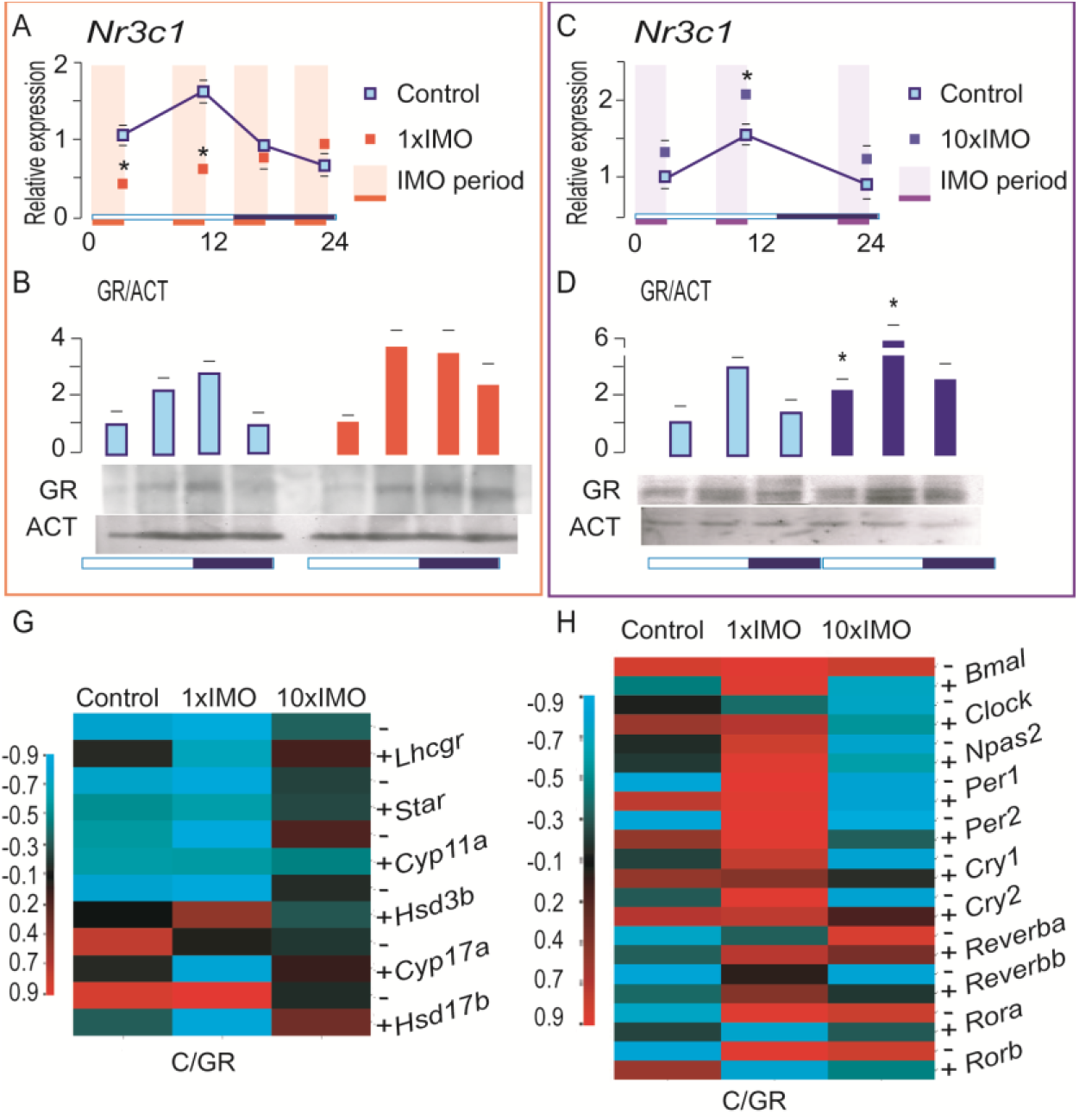
The time-dependent effects of IMO on the expression of the glucocorticoid receptor gene and protein level in Leydig cells. Wistar rats were subject to 1xIMO for three hours in four-time points (ZT0-3, ZT8-11, ZT14-17, ZT20-24) or to 10xIMO in three-time points (ZT0-3, ZT8-11, ZT20-24) during the 24h. Leydig cells were purified, RNA extracted, and used for RT followed by RQ-PCR analysis of *Nr3c1* gene expression (1xIMO: A; 10xIMO: C;). Data points are group mean ± SEM values, (n = 5-6). For possible rhythm prediction please see Suppl. Table 2. The protein level of GR (1xIMO: B; 10xIMO: D;) was determined by western blot. Representative blots are shown. Bars above blots represent mean ± SEM values from pooled data using scanning densitometry values normalized on ACTB. ^*^Statistical significance between Control and IMO group at the same time point (p < 0.05). For The Clustered Image Map datasets show R values from correlation analysis between GR-signaling (presented as a blood corticosterone/GR ratio) and transcription of steroidogenesis-related genes (G) or clock genes (H) during inactive (-) and active (+) phase of the day at different experimental conditions. The red line represents the genes positively correlated with GR-signaling. The blue line corresponds to the genes that were negatively correlated with the GR-signaling.

Since several steroidogenesis-related and clock genes showed different expression patterns regarding the time and type of IMO session, we perform a correlation study to assess the possible association with GR-signaling. The Cluster Image Map (Fig. 4G, H) represents the correlation between the GR-signaling (presented as a ratio of the blood corticosterone/GR in Leydig cells) and the transcriptional level of several steroidogenesis-related (Fig. 4G) and clock genes (Fig. 4H) in Leydig cells. The map presents a relationship of GR-signaling and these two groups of genes in inactive (-) and active (+) phase of the day in the control, 1xIMO or 10xIMO conditions.

In the inactive phase of the day in the control and 1xIMO groups, the analysis showed a negative correlation between C/GR and *Lhcgr, Star, Cyp11a1*, and *Hsd3b* (Fig. 4G). A positive correlation in the control and 1xIMO group’s inactive phase was observed between C/GR and *Hsd17b* (Fig. 4G). In the 10xIMO condition, R values for all investigated genes vary between -0.4 and 0.4 (Fig. 4G). Clock genes, such are *Bmal1, Npas2, Per1,2*, and *Cry1,2*, positively correlate with C/GR in conditions when 1xIMO occurred in both inactive and active phase. Furthermore, a positive correlation was observed for *Rora* and *Rorb* in the inactive phase, and negative in the active phase of the day (Fig. 4H). However, the opposite was registered for *Rev-ereba*, i.e., negative in the inactive and positive in the active phase (Fig. 4H). In the 10xIMO conditions, regardless of the phase of the day, a negative correlation was observed with *Clock, Npas2, Per1,2*. The *Cry1,2*, and *Rev-erbb*, negatively correlate with C/GR in the inactive period, while *Rev-erba, Rora*, and *Rorb* positively (Fig. 4H).

## Discussion

Stress is challenged by any stimulus that disrupts or threatens homeostatic equilibrium. The stress response engages multiple coordinated processes to restore homeostasis and preserve life ^14^. However, reproductive functions are suppressed under various stress conditions depending on genetic predisposition to stress, type, and duration of stress.

Besides that, results from the present study strongly suggest diurnal time as the critical element in determining the stress response. Namely, our previous work pointed to diurnal expression of Leydig cell’s clock genes (*Bmal1, Per1/2/3, Cry1/2, Rev-erba*/*b, Rorb, Dec1/2, Dbp*, and *E4bp4*) but also genes responsible for testosterone production (*Star, Cyp11a1*, and *Cyp17a1*) ^7,8^. However, the expression pattern of these genes changed due to stress and varied depending on the time of the stressful event. In the inactive (light) phase, the critical steroidogenesis-related genes (*Lhcgr, Cyp11a1, Cyp17a1, Hsd3b1/2)* were down-regulated, but in the active (dark) phase of the day, they were unchanged or even up-regulated. The PCA confirmed significant separation of effects on steroidogenesis-related genes depending on the time of the stress in both types of stress used.

It is well known that all steroidogenic events in Leydig cells are initiated and depend on LH-receptor activation, including expression of proteins involved in steroidogenesis such are *Star*/StAR, *Cyp11a1*/CYP11A, *Hsd3b*/HSD3B, *Cyp17a*/CYP17A, and *Hsd17b*/HSD17B (Payne and Hales, 2004). However, results indicate severe inhibitory IMO effects on *Lhcgr* transcription in inactive but not in the active period. This effect did not depend on the type of stress applied. Since LH-receptor signaling is crucial for steroidogenic genes transcription and steroidogenesis activation ^28,29^ decreased expression of *Lhcgr* due to IMO in the rest period could make a difference in respect to IMO in the active period. Also, IMO decreases transcription of pituitary genes encoding LH subunits, *Cga* during the rest and at the beginning of the active period together with *Lhb* in both types of stress. Changes of both genes likely have consequences for the quantity of pulsatile LH secretion and impaired Leydig cells stimulation.

Nevertheless, the IMO is changing the diurnal organization of clocks in the Leydig cells. IMO in both conditions applied potentiates the expression of *Bmal1*/BMAL1 and *Per1/2*/PER1 and in the inactive period decreased *Rev-erba*, suggesting that stressful stimuli can entrain the clock in the Leydig cells. Further, BMAL1 is cyclically expressed in the rat ^6,8^ and mouse ^30^ Leydig cells, stimulating *StAR* expression ^31^ therefore contributing to the cyclic expression of StAR protein. Since IMO in the dark phase promotes BMAL1, this may mitigate the effects of stress on steroidogenesis during the night. Also, the promoters of steroidogenic genes such are *Star, Cyp11a1*, and *Cyp17a1* ^30^ as well as promoter of steroidogenic regulators such as *Nur77* contain clock-controlled RORE and E-box, suggesting BMAL1, PERs, and REV-ERBs as potential regulators of rhythmic expression of steroidogenic genes. Further studies are necessary to confirm this hypothesis.

The clock system interacts with many different signals to produce an integrated output over the 24h cycle, entraining other cyclic activities in the cell ^32^. It is well known that rhythmic corticosterone secretion is an important timing signal for the coordination of the peripheral clock ^33^. Both circadian and stress-mediated aspects play a crucial role in determining the level of corticosterone secretion and mediating the effects on target cells. Corticosterone acts as a peripheral clock synchronizer ^34,35^ by regulation of clock genes expression through genomic actions mediated by activation of GR ^36^. Actually, it was found that *Per1* and *Per2* contain glucocorticoid-responsive elements. Further, *Rev-erba* and *Rora* are negatively regulated by glucocorticoids ^37,38^. Results from this study showed altered *Nr3c1*/GR expression profile in Leydig cells due to IMO. Thus, results suggest that the diurnal effect of stress is mediated, among others, by the circadian dynamics of corticosterone signaling.

Indeed, correlation study suggested a positive relationship between GR signaling (C/GR) and transcriptional activity of the clock *Bmal1, Per1,2, Cry1,2*, and *Rora,b* and negative with steroidogenesis-related *Lhcgr, Star, Cyp11a1*, and *Hsd3b* when 1xIMO occurred in the inactive phase of the day. However, when IMO repeatedly occurred in the inactive period, a negative relationship with *Clock, Npas2, Per1,2, Cry1,2*, and *Rev-erbb* suggested different regulation in 1xIMO and repeated IMO conditions. Additionally, the diurnal pattern of the *Nr3c1*/GR expression in Leydig cells was changed ie increased in the 10xIMO, which contributes to the different effect observed. However, many other signaling pathways directed by hormones, especially LH, PRL, adrenaline, involved in highly multi-regulated processes such as gene expression, could be affected by IMO and vary the circadian rhythm.

In summary, the results suggest different sensitivity to stress depending on the diurnal time of the stressful event. Transcriptions of the clock and steroidogenic genes are much more affected by stress in the rest than in the active phase, pointed to the relevance of the stress event’s timing. Both types of stress (1xIMO and 10xIMO) potentiates expression of clock elements that act over E-box (BMAL1 and PER1) regardless of the phase of the day and decreased elements which exert their action via RORE-box (*Rev-erba,b*) in rest period, suggesting a possible effect on gene expression not only on the clock system but also outside it.

The obtained results pointed to stress and the circadian system’s coordinated action in shaping male reproductive physiology, emphasizing the importance of the Leydig cells’ rhythmic activity underpinning male fertility.

## Author Contributions

M.LJ.M.— data acquisition; data analysis and interpretation; figures drafting; revising manuscript critically for important intellectual content; S.A.A.—acquisition of the data; analysis and interpretation of the data; revising manuscript critically for important intellectual content; T.S.K.—the conception and design of the research; acquisition of the data; analysis and interpretation of the data; drafting the manuscript; All authors—approved the final version of the manuscript; agree to be accountable for all aspects of the work in ensuring that questions related to the accuracy or integrity of any part of the work are appropriately investigated and resolved.

## Acknowledgments

We are very grateful to Professor dr Gordon Niswender (Colorado State University) for supplying antibodies for radioimmunoassay analysis. Also we thank dr Aleksandar Baburski for technical assistance including data analysis as well as dr Milan Župunski for technical help in drafting figures.

