## Supplemental Tables 1-8 for "Stress alters the activity of Leydig cells dependently on the diurnal time"

**Supplemental Table 1.:** The primers sequences used for real time PCR analysis.

| **Gene** | **GenBank accession code** | **Primers sequences** | |
|---|---|---|---|
| *Gapdh* | NM_017008 | F:5'-TGCCAAGTATGATGACATCAAGAAG-3' | R:5'-AGCCCAGGATGCCCTTTAGT-3' |
| *Cga* | NM_053918.2 | F:5'-CAGTGTATGGGCTGTTGCTTCT-3' | R:5'-GGAACCAACATTGTCTTCTTGGA-3' |
| *Lhb* | NM_012858.2 | F:'TCTTCTGATGCCCACCCACTA-3' | R:5'-TATTGGGAGGGATGGTTAGAACA-3' |
| *Lhr* | NM_012978 | F:5'-CGGGCTGGAGTCCATTCA-3' | R:5'-TTCTTTGGAGGGCAGTGTTTTC-3' |
| *Nr3c1* | NC_005117.4 | F:5'-CGGTTAATCTGCACAGCCTAT -3' | R:5'-AAAATGGGTCGGTGCTTCTA -3' |
| *Star* | NM_031558 | F:5'-AGCCAGCAGGAGAATGGAGAT-3' | R:5'-CACCTCCAGTCGGAACACCTT-3' |
| *Cyp11a1* | NM_017286 | F:5'-GGGCAACATGGAGTCAGTTTACA-3' | R:5'-GACCCTCGCAGGAGAAGAGA-3' |
| *Hsd3b1/2* | NM_001042619.1 | F:5'-GACAGGAGCAGGAGGGTTTGTGG-3' | R:5'-CTCCTTCTAACATTGTCACCTTGGCCT-3' |
| *Cyp17a1* | NM_012753 | F:5'-GCCACGGGCGACAGAA-3' | R:5'-GCCTTTGTTGGGAAAAATCG-3' |
| *Hsd17b4* | NM_024392 | F:5'-CCTTTGGCTTTGCCATGAGA-3' | R:5'-CAATCCATCCTGCTCCAACCT-3' |
| *Clock* | NM_021856.1 | F:5'- ACAGCCCCACTGTACAATACGA-3' | R:5'- TGCGGCATACTGGATGGAAT-3' |
| *Bmal1* | NM_024362.2 | F:5'- AAGAGGCGTCGGGACAAAAT-3' | R:5'- TTCCGGGACATCGCATTG-3' |
| *Npas2* | NM_001108214.2 | F:5'- GGCGCACCCTGTGTACATTT-3' | R:5'- TCTTTCCCCATTCTGCAAGTG-3' |
| *Per1* | NM_001034125.1 | F:5'- CCTGCACACCCAGAAGGAA-3' | R:5'- GAGGTGTCAAGCCCACGAA-3' |
| *Per2* | NM_031678.1 | F:5'- GGAAGGAGGCCCAGACGTA-3' | R:5'- TGGGTCCATTTCGTTAGAAACA-3' |
| *Cry1* | NM_198750.2 | F:5'- ACCATCCGCTGCGTGTACAT-3' | R:5'- AGCAAAAATCGCCACCTGTT-3' |
| *Cry2* | NM_133405.1 | F:5'- TTCCCAAGGCTTTTCAAGGA-3' | R:5'- TCCCGTTCTTTCCCAAAGG-3' |
| *Rora* | NM_001106834.1 | F:5'- GAAGAACCACCGAGAAGATGGA-3' | R:5'- CGTCCGCATAGGGCTCTTAA-3' |
| *Rorb* | NM_001270958.1 | F:5'- CAGGAACCGTTGCCAACAC-3' | R:5'- GGACATCCTCCCAAACTTCACA -3' |
| *Rev-erba* | NC_005109 | F:5'- GAGCATCCAGCAGAACATCCA-3' | R:5'- TTGCGATTGATACGGACAATG-3' |
| *Rev-erbb* | NC_005114.4 | F:5'- GAACGAGAATTGCTCCATCATG -3' | R:5'- CGACATTCCCACGGACAGA -3' |

Primers were design by using Primer Express 3.0 software (Applied Biosystems) and full genes sequences from NCBI GenBank (http://www.ncbi.nlm.nih.gov/nuccore). F - forward, R – reverse.

**Supplemental Table 2. Rhythm parameters for RQ-PCR analysis of gene expression present in Fig.2 and Fig.3.**

| | Group | p | Mesor | Amplitude | Acrophase |
|---|---|---|---|---|---|
| **Testosterone** | **Control** | 0.000010 | 4.126340 | 2.709868 | 10.392056 |
| **Corticosterone** | **Control** | 0.00001 | 57.379546 | 28.256135 | 9.807214 |
| **Glucose** | **Control** | 0.053450 | 6.991290 | 0.506288 | 9.121611 |
| ***Cga*** | **Control** | 0.000010 | 2.539760 | 1.344414 | 15.387778 |
| ***Cgb*** | **Control** | 0.000010 | 1.169380 | 0.414669 | 18.451450 |
| ***Lhcgr*** | **Control** | 0.080400 | | | |
| ***Nr3c1*** | **Control** | 0.000050 | 1.103210 | 0.325498 | 10.376503 |
| ***Star*** | **Control** | 0.040120 | 0.996430 | 0.295860 | 3.168410 |
| ***Cyp11a1*** | **Control** | 0.000120 | 0.801560 | 0.394170 | 6.150374 |
| ***Hsd3b1/2*** | **Control** | 0.143420 | | | |
| ***Hsd17b4*** | **Control** | 0.000005 | 0.830570 | 0.115314 | 2.849319 |
| ***Cyp17a1*** | **Control** | 0.000050 | 0.705630 | 0.347059 | 6.061222 |
| ***Bmal1*** | **Control** | 0.000001 | 0.560690 | 0.408230 | 1.909399 |
| ***Clock*** | **Control** | 0.361930 | | | |
| ***Npas2*** | **Control** | 0.060000 | | | |
| ***Per1*** | **Control** | 0.000190 | 1.165 | 0.5610238 | 11.587508 |
| ***Per2*** | **Control** | 0.000010 | 1.802710 | 0.881767 | 12.759931 |
| ***Cry1*** | **Control** | 0.000050 | 1.868170 | 1.461789 | 18.230596 |
| ***Cry2*** | **Control** | 0.00805 | 0.748620 | 0.350629 | 9.164588 |
| ***Rev-erba*** | **Control** | 0.000960 | 1.005010 | 0.334595 | 7.523166 |
| ***Rev-erbb*** | **Control** | 0.000001 | 0.885530 | 0.659719 | 7.522000 |
| ***Rora*** | **Control** | 0.585500 | | | |
| ***Rorb*** | **Control** | 0.000150 | 0.795850 | 0.795850 | 8.297699 |

**Supplemental Table 3.** Variables loadings from principal component analysis. PC-principal component.

| | PC1 | PC2 | PC3 |
|---|---|---|---|
| Lhgcr | -0.07152 | 0.491352 | 0.642671 |
| Star | -0.50755 | -0.39841 | 0.235081 |
| Cyp11 | -0.52355 | -0.09524 | -0.20899 |
| Cyp17a1 | 0.080507 | -0.59493 | 0.623839 |
| Hsd3b1.2 | -0.57174 | -0.01094 | -0.17463 |
| Hsd17b | 0.36029 | -0.48653 | -0.26146 |

**Supplemental Table 4.** Variables loadings from principal component analysis. PC-principal component.

| | PC1 | PC2 | PC3 |
|---|---|---|---|
| Lhgcr | 0.52014 | -0.20273 | -0.15335 |
| Star | 0.335387 | 0.467236 | 0.572887 |
| Cyp11a1 | 0.483617 | 0.024252 | 0.053696 |
| Cyp17a1 | -0.02927 | 0.84484 | -0.47888 |
| Hsd3b1.2 | 0.451647 | 0.057046 | 0.227036 |
| Hsd17.b4 | 0.422188 | -0.15164 | -0.60377 |

**Supplemental Table 5.** Variables loadings from principal component analysis. PC-principal component.

| | PC1 | PC2 | PC3 |
|---|---|---|---|
| Lhgcr | -0.4523 | -0.0326 | -0.0254 |
| Star | -0.4271 | -0.3530 | -0.2330 |
| Hsd3b1.2 | -0.4556 | -0.0336 | -0.0369 |
| Cyp11a1 | -0.4482 | 0.0260 | -0.4677 |
| Cyp17a1 | -0.1510 | 0.9317 | -0.1378 |
| Hsd17b1.2 | 0.4264 | -0.0669 | -0.8402 |

**Supplemental Table 6.** Variables loadings from principal component analysis. PC-principal component.

| | PC1 | PC2 | PC3 |
|---|---|---|---|
| Clock | -0.25404 | 0.068451 | -0.49355 |
| Npas2 | -0.32152 | 0.206485 | 0.186631 |
| Bmal1 | 0.123285 | 0.313721 | -0.56317 |
| Rora | -0.10154 | -0.51169 | -0.1372 |
| Rorb | -0.44544 | -0.06615 | -0.13804 |
| Cry1 | 0.297963 | 0.188317 | 0.431501 |
| Cry2 | -0.36158 | 0.360493 | -0.05679 |
| Per1 | -0.39623 | 0.214477 | 0.189347 |
| Per2 | -0.35392 | 0.040101 | 0.373339 |
| Rev.erba | -0.03673 | -0.48991 | 0.019381 |
| Rev.erbb | -0.3241 | -0.36745 | 0.038873 |

**Supplemental Table 7.** Variables loadings from principal component analysis. PC-principal component.

| | PC1 | PC2 | PC3 |
|---|---|---|---|
| Clock | 0.29394 | 0.252515 | -0.012 |
| Npas2 | 0.418848 | 0.12091 | -0.16573 |
| Bmal1 | -0.01235 | -0.26517 | 0.64281 |
| Rora | 0.291554 | 0.350113 | -0.30549 |
| Rorb | -0.16715 | 0.517165 | 0.036471 |
| Cry1 | -0.28311 | -0.39929 | -0.19258 |
| Cry2 | -0.34858 | 0.323843 | 0.132776 |
| Per1 | -0.39162 | 0.287647 | 0.001399 |
| Per2 | -0.21037 | -0.22199 | -0.58538 |
| Rev.erba | -0.30645 | -0.04314 | -0.24555 |
| Rev.erbb | -0.36315 | 0.251142 | 0.082741 |

**Supplemental Table 8.** Variables loadings from principal component analysis. PC-principal component.

| | PC1 | PC2 | PC3 |
|---|---|---|---|
| Clock | -0.17029 | 0.417045 | 0.266498 |
| Npas2 | 0.360635 | -0.2167 | -0.15463 |
| Bmal1 | 0.360635 | -0.2167 | -0.15463 |
| Rora | 0.272342 | 0.353571 | -0.06742 |
| Rorb | 0.407769 | 0.001224 | -0.04669 |
| Cry1 | -0.39712 | -0.03046 | 0.123363 |
| Cry2 | -0.26365 | -0.34202 | -0.0533 |
| Per1 | 0.151323 | -0.35597 | 0.881661 |
| Per2 | -0.36249 | 0.201738 | -0.05567 |
| Rev.erba | 0.264825 | 0.358734 | 0.175803 |
| Rev.erbb | -0.13708 | -0.43948 | -0.21223 |
